# The two chemotaxis in *Caulobacter crescentus* operons play different roles in chemotaxis and biofilm regulation

**DOI:** 10.1101/528224

**Authors:** Cécile Berne, Yves V. Brun

## Abstract

The holdfast polysaccharide adhesin is crucial for irreversible cell adhesion and biofilm formation in *Caulobacter crescentus*. Holdfast production is tightly controlled via developmental regulators, and environmental and physical signals. Here we identified a novel mechanism of holdfast production regulation that involves chemotaxis proteins. We characterized the two identified chemotaxis operons of *C. crescentus* and showed that only the previously characterized, major operon is involved in chemotactic response towards different carbon sources. However, both chemotaxis operons encoded in the *C. crescentus* genome play a role in biofilm formation and holdfast production, by regulating the expression of *hfiA*, the gene encoding the holdfast inhibitor HfiA. We show that CheA and CheB proteins act in an antagonistic manner: while the two CheA proteins negatively regulate *hfiA* expression, the CheB proteins are positive regulators, thus providing a modulation of holdfast synthesis and surface attachment.

**IMPORTANCE:** Chemosensory pathways are major signal transduction mechanisms in bacteria. These systems are involved in chemotaxis and other cell responses to environment conditions, such as production of adhesins that enable irreversible adhesion to a surface and surface colonization. The *C. crescentus* genome encodes two complete chemotaxis operons. Here we characterized the second, novel chemotaxis-like operon. While only the major chemotaxis operon is involved in chemotaxis, both chemotaxis systems modulate *C. crescentus* adhesion by controlling expression of the holdfast synthesis inhibitor, HfiA. Thus, we identified a new level in holdfast regulation, providing new insights into the control of adhesin production that leads to the formation of biofilms.

## INTRODUCTION

In their natural habitat, most bacteria are organized in complex surface-associated, multicellular communities known as biofilms. The first step of biofilm formation is the reversible adhesion of a few single cells to a surface. When conditions are favorable, these attached cells produce adhesin molecules, which strengthen the interaction with the surface. The cells then divide to form multicellular microcolonies, which eventually develop into a mature biofilm (1). Communal life on a surface is believed to be beneficial, as it provides protection from predators and xenobiotic stresses (2). The environment at the surface is highly heterogenous, with the presence of various compounds adsorbed on the surface and the formation of gradients near it (1). To initiate attachment, bacteria must approach the surface either by passive transport or by active swimming (1). Active swimming toward the surface and initial surface attachment can be biased by environmental cues and chemotaxis (3, 4). For example, chemotaxis is involved in the colonization of biotic (5, 6) and abiotic surfaces (7–9) and in cell-cell aggregation (10, 11). Finally, chemotaxis is also involved in later stages of biofilm formation, as is the case for *s*ingle *Pseudomonas aeruginosa* cells within a mature biofilm that can actively respond to a chemical gradient and subsequently reposition themselves on the surface (12).

The central chemotaxis system is composed of a chemoreceptor, or <u>m</u>ethyl-accepting <u>c</u>hemotaxis <u>p</u>rotein (MCP), and CheW, CheA, and CheY proteins (13–16). Complexes of these proteins form hexagonally packed arrays localized at the cell pole. MCPs are transmembrane proteins, while CheW, CheA, and CheY are located in the cytoplasm (17, 18). Chemotactic signals are sensed by the MCPs and transduced to the sensory histidine kinase CheA, which is linked to the MCP via the scaffolding protein CheW (Fig. 1A). CheA autophosporylates upon receiving the signal and rapidly transfers its phosphate to the CheY response regulator, which modulates flagellum rotation. CheA can also transmit its phosphate to the methylesterase CheB, at a slower rate than the transfer to CheY. CheB~P is activated and removes a methyl group from the MCP, reducing the activity of the signal transduction cascade to control the MCP adaptation process that allows the signal to be reset to a background state so bacteria can detect and respond to an increasing concentration gradient of signal. Optional accessory proteins have also been linked to the chemotaxis apparatus, such as CheC and its homolog CheX, CheD, CheV, CheE, and CheZ (13).

**Figure 1:**
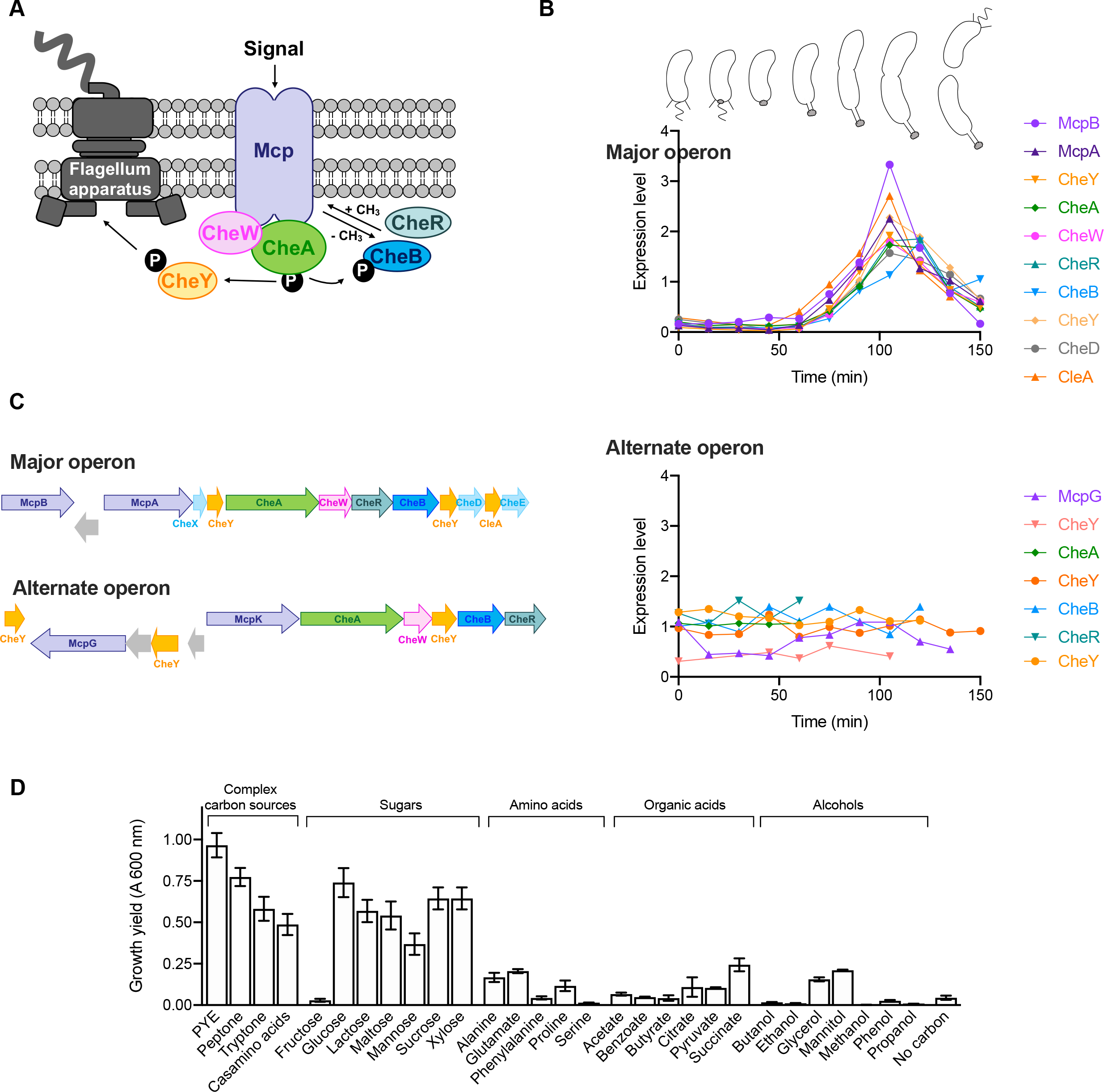
Chemotaxis operon organization in *C. crescentus* and growth using different carbon sources. **(A)** Schematic representation of the central chemotaxis apparatus. **(B)** Cell-cycle-dependent transcriptional regulation of genes encoded in the major and alternate chemotaxis operons. Data were extracted from previous global transcriptome analysis (19). Approximate *C. crescentus* cell-cycle progression is shown on top. **(C)** Genomic organization of the major and alternate chemotaxis operons (gene accession numbers are given in Table 1). Genes encoding the central chemotaxis apparatus are represented using the same colors as in Fig. 1A; chemotaxis accessory proteins are represented in cyan **(D)** Growth yield (OD_600_ after 24 h of incubation at 30°C) of CB15 WT grown using different carbon sources. Results are given as the mean of 3 independent replicates, and error bars represent the standard error of the mean (SEM).

In *Caulobacter crescentus*, the chemotaxis apparatus forms concomitantly with the flagellum apparatus (19, 20). This dimorphic bacterium starts its life as a piliated, flagellated, motile swarmer cell and then transitions to a sessile, stalked cell by retracting its pili, shedding its flagellum, and synthesizing an adhesive holdfast followed by a stalk at the same pole (Fig. 1B). The chemoreceptor McpA is the best characterized MCP in *C. crescentus*, and it is synthesized in a cell-cycle dependent manner, similarly to the other chemotaxis proteins encoded in this major chemotaxis operon (19, 21–24) (Fig. 1B). McpA is synthesized at the new pole of predivisonal cells; thus, newborn swarmer cells inherit McpA at the flagellar pole after division (25, 26). McpA is then degraded during the swarmer to stalked-cell transition (27) via proteolysis by ClpX (28), and this temporally regulated proteolysis plays an important role in the asymmetric distribution of McpA (27).

**Table 1:** Putative chemotaxis genes of *C. crescentus*.

| Gene symbol | Locus tag in CB15 | Locus tag in NA1000 | Gene product |
| --- | --- | --- | --- |
| **Major chemotaxis operon** |  |  |  |
| *mcpB* | CC_0428 | CCNA_00437 | Receptor methyl-accepting chemotaxis protein MCP |
| *mcpA* | CC_0430 | CCNA_00439 | Receptor methyl-accepting chemotaxis protein MCP |
| *cheY* | CC_0432 | CCNA_00441 | Response regulator protein CheY |
| *cheA* | CC_0433 | CCNA_00442 | Chemotaxis sensory histidine kinase CheA |
| *cheW* | CC_0434 | CCNA_00443 | Receptor binding protein CheW |
| *cheR* | CC_0435 | CCNA_00444 | Chemotaxis protein methyltransferase CheR |
| *cheB* | CC_0436 | CCNA_00445 | Chemotaxis protein methyltransferase CheB |
| *cheY* | CC_0437 | CCNA_00446 | Response regulator protein CheY |
| *cheD* | CC_0438 | CCNA_00447 | Chemotaxis protein CheD |
| *cleA* | CC_0440 | CCNA_00449 | CheY-like cdG effector CleA |
| **Alternate chemotaxis operon** |  |  |  |
| *cheY* | CC_0588 | CCNA_00625 | Response regulator protein CheY |
| *mcpG* | CC_0589 | CCNA_00626 | Receptor methyl-accepting chemotaxis protein MCP |
| *cheY* | CC_0591 | CCNA_00628 | Response regulator protein CheY |
| *mcpK* | CC_0593 | CCNA_00629 | Receptor methyl-accepting chemotaxis protein MCP |
| *cheA* | CC_0594 | CCNA_00630 | Chemotaxis sensory histidine kinase CheA |
| *cheW* | CC_0595 | CCNA_00631 | Receptor binding protein CheW |
| *cheY* | CC_0596 | CCNA_00632 | Response regulator protein CheY |
| *cheB* | CC_0597 | CCNA_00633 | Chemotaxis protein methyltransferase CheB |
| *cheR* | CC_0598 | CCNA_00634 | Chemotaxis protein methyltransferase CheR |
| **Single genes** |  |  |  |
| *mcpQ* | CC_0066 | CCNA_00064 | Receptor methyl-accepting chemotaxis protein MCP |
| *mcpC* | CC_0343 | CCNA_00160 | Receptor methyl-accepting chemotaxis protein MCP |
| *mcpN* | CC_0504 | CCNA_00538 | Receptor methyl-accepting chemotaxis protein MCP |
| *cheW* | CC_0764 | CCNA_00803 | Receptor binding protein CheW |
| *cleB* | CC_1364 | CCNA_01426 | CheY-like cdG effector CleB |
| *mcpP* | CC_1399 | CCNA_01465 | Receptor methyl-accepting chemotaxis protein MCP |
| *mcpD* | CC_1655 | CCNA_01727 | Receptor methyl-accepting chemotaxis protein MCP |
| *cleC* | CC_2249 | CCNA_02332 | CheY-like cdG effector CleC |
| *mcpE* | CC_2281 | CCNA_02364 | Receptor methyl-accepting chemotaxis protein MCP |
| *mcpM* | CC_2317 | CCNA_02402 | Receptor methyl-accepting chemotaxis protein MCP |
| *mcpF* | CC_2691 | CCNA_02773 | Receptor methyl-accepting chemotaxis protein MCP |
| *mcp* | CC_2810 | CCNA_02901 | Receptor methyl-accepting chemotaxis protein MCP |
| *mcpO* | CC_2842 | CCNA_02773 | Receptor methyl-accepting chemotaxis protein MCP |
| *mcpI* | CC_2847 | CCNA_02901 | Receptor methyl-accepting chemotaxis protein MCP |
| *cheW* | CC_3025 | CCNA_03120 | Receptor binding protein CheW |
| *cleD* | CC_3100 | CCNA_03198 | CheY-like cdG effector CleD |
| *mcpJ* | CC_3145 | CCNA_03247 | Receptor methyl-accepting chemotaxis protein MCP |
| *cleE* | CC_3155 | CCNA_03257 | CheY-like cdG effector CleE |
| *cheY* | CC_3258 | CCNA_03367 | Response regulator protein CheY |
| *mcpH* | CC_3349 | CCNA_03459 | Receptor methyl-accepting chemotaxis protein MCP |
| *cheY* | CC_3471 | CCNA_03585 | Response regulator protein CheY |
| *cheR* | CC_3472 | CCNA_03586 | Chemotaxis protein methyltransferase CheR |

*C. crescentus* irreversibly adheres to surfaces and forms a biofilm by producing an adhesive holdfast (29–32). This polysaccharide adhesin is composed of β-1,4 *N*-acetyl-glucosamine residues (33, 34), peptides, and DNA molecules (35). In complex medium, holdfast synthesis is temporally regulated via two pathways: a developmental program, or contact with a surface (1, 32). Newborn *C. crescentus* cells spend approximately one-third of their lifespan as motile swarmer cells before differentiating though highly controlled cell-cycle progression into cells synthesizing holdfast and a stalk (Fig. 1B). However, swarmer cells that reach a surface bypass this developmental pathway and synthesize a holdfast within seconds of surface contact (36–39). The genes involved in holdfast synthesis and anchoring are transcribed in predivisional cells, resulting in newborn swarmer cells bearing complete and functional holdfast machinery. Holdfast production is regulated post-transcriptionally. The second messenger molecule cyclic-di-GMP (cdG) is the main regulator of holdfast production; levels of intracellular cdG increase during the swarmer to stalked cell differentiation, triggering the transition from motile to sessile states by flagellum shedding and holdfast production (40–42). In addition, HfsJ, a glycosyltransferase required for holdfast synthesis, directly binds cdG, triggering holdfast synthesis upon contact with the surface (39). The holdfast inhibitor HfiA also regulates holdfast synthesis by inhibiting HfsJ (43). The transcription of *hfiA* is cdG dependent (44) and cell-cycle regulated, with transcript production occurring in stalked and predivisional cells and transcripts being degraded in swarmer cells (43). Environmental factors, such as blue light via the LovK-LovR system and nutrient availability, add additional control of *hfiA* expression (43, 45). Finally, HfiA is also regulated posttranscriptionally by interaction with the chaperone DnaK, ensuring its stabilization in the cell (46).Overall, this multilayered control ensures that holdfast production and subsequent irreversible adhesion are tightly controlled at different levels.

In this study, we investigated the role of chemotaxis in biofilm formation and holdfast production under different nutrient conditions in *C. crescentus*. We focused on dissecting the role of both the major and alternate chemotaxis operons in motility and adhesion. While previous works largely focus on the role of genes encoded by the major chemotaxis operon in swimming behaviors, here we analyzed the role of the second, alternate chemotaxis operon by comparing mutants lacking the key CheA-type response regulators and the CheB-type methyltransferases that are encoded in the two chemotaxis operons. We first show that only the major chemotaxis operon is involved in chemotaxis, as only *ΔcheAI* and *ΔcheBI* mutants in the major operon were unable to respond to a chemotactic gradient, while *ΔcheAII* and *ΔcheBII* mutants in the alternate operon behaved like WT. We then demonstrate that both operons play a role in cell attachment and holdfast production in a complex, nutrient-dependent manner. CheA and CheB proteins act antagonistically: CheAI and CheAII positively regulate adhesion, while CheBI and CheBII repress it. These proteins also act to control the expression of the gene encoding the holdfast inhibitor, HfiA. These results highlight different roles in regulating chemotaxis and biofilm formation for the two chemotaxis operons.

## RESULTS

### Genomic organization of the chemotaxis genes in *C. crescentus*

There are two chemotaxis operons encoded in the *C. crescentus* genome (47, 48). Historically, all mutations impairing the chemotactic response have been identified in a single operon (26, 27, 49, 50), referred to as the major chemotaxis operon (28). Hence, we named the second operon the alternate chemotaxis operon. While the major operon is cell-cycle regulated (19, 20, 23, 24), with a peak of expression occurring in predivisional cells, the alternate operon was constitutively expressed during the cell-cycle (Fig. 1B).

Both operons are arranged similarly, with two MCPs, one CheA histidine kinase, three CheY response regulators, one CheW, one CheR, and one CheB (Fig. 1C). While the major operon encodes a copy of CheD and CheE accessory proteins, both are absent from the alternate operon. In addition to the four MCP-encoding genes present in the two chemotaxis operons, there are also 14 independent genes coding for putative MCPs (Table 1), suggesting that *C. crescentus* may sense a large array of specific attractants. The presence of a large number of MCPs in its genome is consistent with *Caulobacter* being a bacterium abundantly found in oligotrophic freshwater and nutrient-rich soil environments (51, 52). There are also several copies of key chemotaxis genes scattered in the genome, such as six copies of *cheY*, two of *cheW*, and one extra *cheR* (Table 1). No *cheC*, *cheV*, or *cheZ* homologs were found in the genome (Table 1). Among the 12 homologs of *cheY*, five have been recently characterized as encoding <u>C</u>heY-<u>l</u>ike cdG <u>e</u>ffectors (Cle) proteins (53). CleA is part of the major chemotaxis operon, while CleB, CleC, CleD and CleE are located independently in the genome (53) (Table 1). These Cle proteins bind to cdG and are involved in tuning of the flagellar motor activity, resulting in the subsequent increase of holdfast production upon contact with the surface (53).

In this study, we investigated the behaviors of in-frame deletions mutants lacking either *cheA*, encoding the central histidine kinase that transduces the signal from the MCP receptor to the key response regulator proteins CheY, or *cheB*, encoding the methyltransferase response regulator that is involved in transferring a methyl group to the receptor MCP and thus modulating the chemotatic response. Genes encoded in the major operon have been previously named *cheAI* and *cheBI* (54), so we named the genes encoded in the alternate operon *cheAII* and *cheBII*, and we constructed in-frame single deletions of each of the *cheA* (*cheAI* and *cheAII*) and *cheB* (*cheBI* and *cheBII*) genes in *C. crescentus* CB15.

### Determination of carbon sources metabolized by *C. crescentus*

To determine the appropriate carbon sources to use in this study, we first surveyed a wide array of compounds that could be metabolized by *C. crescentus* as the sole carbon source by testing a range of complex carbon sources, sugars, amino acids, organic acids, and alcohols. We monitored the growth yield obtained in M2 medium supplemented with a single given carbon source after 24 hours and compared it to the growth yield and generation time in complex <u>p</u>eptone-<u>y</u>east <u>e</u>xtract (PYE) medium (Fig. 1D and Table 2). While M2 defined media provide inorganic phosphate, ammonium salts, and carbon, bacteria must *de novo* synthesize amino acids and nucleotides, which are crucial for growth. *C. crescentus* preferentially grew using complex carbon and sugars (Fig. 1D), confirming previous observations (51, 55). Mutations in the major and alternate chemotaxis operons did not impair growth (Table 2 and Fig. S1). We decided to focus on carbon sources metabolized more efficiently by *C. crescentus* and chose tryptone as an example of complex carbon source, and glucose, maltose, sucrose, and xylose as examples of sugars, to conduct our studies. Interestingly, with the exception of sucrose, these sugars have been previously reported to be specific chemoattractant sugars for *C. crescentus* (49, 56, 57).

**Table 2:** Generation times (in minutes) of WT and mutant strains of *C. crescentus* CB15 grown using different carbon sources.

| | WT | $\Delta cheA$I | $\Delta cheB$I | $\Delta cheA$II | $\Delta cheB$II |
| --- | --- | --- | --- | --- | --- |
| **Complex carbon sources** |  |  |  |  |  |
| PYE | 94 ± 4.5 | 95 ± 5.1 | 97 ± 4.6 | 93 ± 1.6 | 95 ± 8.0 |
| Casamino acids | 123 ± 4.5 | 122 ± 7.2 | 116 ± 8.3 | 125 ± 5.2 | 119 ± 6.8 |
| Peptone | 115 ± 5.3 | 128 ± 3.9 | 110 ± 5.5 | 118 ± 3.7 | 125 ± 7.9 |
| Tryptone | 109 ± 2.8 | 102 ± 4.8 | 115 ± 6.7 | 113 ± 6.2 | 105 ± 6.6 |
| **Sugars** |  |  |  |  |  |
| Glucose | 121 ± 5.9 | 125 ± 7.6 | 122 ± 8.2 | 118 ± 4.9 | 120 ± 8.2 |
| Maltose | 125 ± 9.4 | 127 ± 6.7 | 132 ± 7.5 | 129 ± 7.2 | 131 ± 8.8 |
| Sucrose | 137 ± 3.3 | 141 ± 5.9 | 144 ± 6.2 | 140 ± 1.4 | 138 ± 5.0 |
| Xylose | 138 ± 7.1 | 138 ± 5.5 | 138 ± 5.4 | 134 ± 5.7 | 135 ± 6.3 |
| **Amino acids** |  |  |  |  |  |
| Alanine | 473 ± 9.8 | 496 ± 10.2 | 487 ± 8.8 | 500 ± 10.1 | 478 ± 9.1 |
| Glutamic acid | 443 ± 8.3 | 437 ± 9.3 | 438 ± 6.9 | 449 ± 8.9 | 444 ± 10.1 |
| **Organic acids** |  |  |  |  |  |
| Citrate | 573 ± 16.8 | 590 ± 9.1 | 578 ± 14.3 | 582 ± 16.2 | 585 ± 8.2 |
| Pyruvate | 569 ± 8.2 | 582 ± 14.2 | 558 ± 9.8 | 580 ± 10.9 | 560 ± 13.1 |
| Succinate | 391 ± 10.9 | 405 ± 6.9 | 401 ± 10.3 | 406 ± 11.7 | 403 ± 10.0 |
| **Alcohols** |  |  |  |  |  |
| Glycerol | 604 ± 11.0 | 613 ± 12.9 | 628 ± 14.0 | 655 ± 12.7 | 599 ± 14.9 |
| Mannitol | 630 ± 12.6 | 623 ± 16.2 | 625 ± 11.4 | 634 ± 8.9 | 622 ± 11.7 |

### Only the major chemotaxis operon is involved in chemotactic response toward different carbon sources

In *C. crescentus*, previous studies on chemotaxis almost exclusively focused on proteins encoded by the major chemotaxis operon, with CheAI (CC_0433) (54, 56), CheBI (CC_0436) (49, 54), and CheRI (CC_0435) (49, 54) receiving most of the attention. A more recent work investigated the chemotactic behavior of CheB and CheR null mutants (in-frame deletions of *cheBI* and *cheBII*, or *cheRI*, *cheRII*, and *cheRIII*) (57), but, to our knowledge, the putative role of the alternate chemotaxis operon in chemotaxis remains unstudied.

We monitored the swimming of *C. crescentus* CB15 wild-type (WT) and mutant strains in semisolid agar plates containing different sugars as sole carbon sources (Fig. 2), as only motile bacteria able to respond to a chemotactic gradient can form a ring under such conditions (49). Mutants in the alternate chemotaxis operon (Δ*cheAII* and Δ*cheBII*) exhibited similar swimming behaviors as WT, suggesting that the alternate operon is not involved in chemotaxis (Fig. 2). However, both mutants in the major chemotaxis operon (Δ*cheAI* and Δ*cheBI*) were impaired in chemotaxis, as deduced from their reduced ability to swim in semisolid media compared to WT (Fig. 2). These phenotypes could be complemented *in trans* by a replicating plasmid encoding a copy of *cheAI* or *cheBI* under control of a constitutive promoter (Fig. S2A).

**Figure 2:**
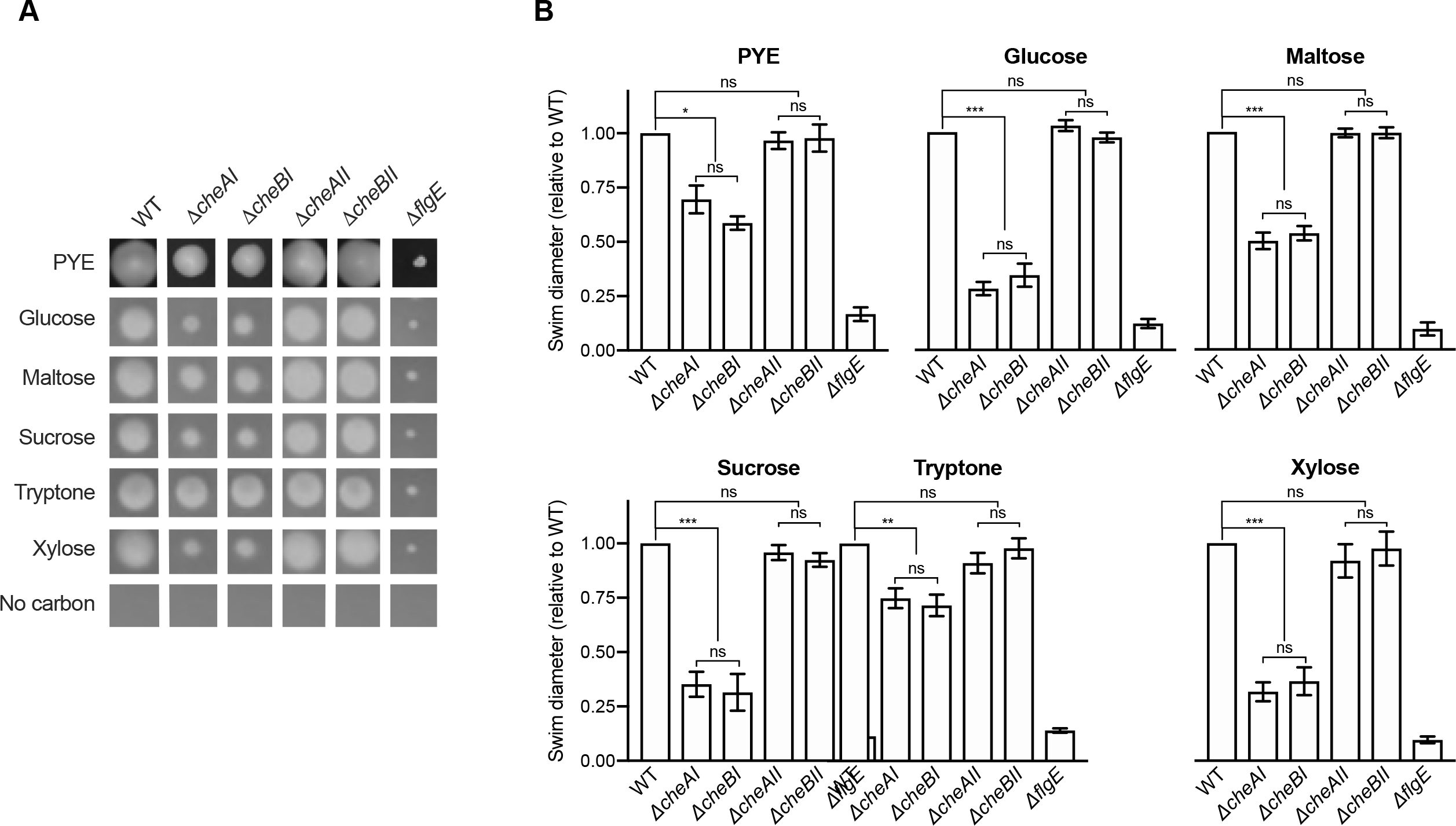
Motility assays in semisolid agar. **(A)** Representative images of swim rings obtained after 5 days of incubation in semisolid plates made with M2 medium + carbon source or PYE + noble agar (0.4%). (B) Swim diameters of the different strains using different carbon sources. Results are normalized to WT ring diameter on the same plate type. Bar graphs indicate the mean of five independent replicates, and error bars represent SEM. Statistical comparisons were calculated using paired t-tests. *** *P* < 0.001; ** *P* < 0.01; * *P* < 0.1; ns = not significant.

The size of the swim ring, and therefore the amplitude of the chemotactic response, was different depending on the carbon source (Fig. 2); there was a 75% decrease for the Δ*cheAI* and Δ*cheBI* mutants when glucose, sucrose, or xylose was used as the carbon source, while a less drastic decrease was observed in the presence of other carbon sources (50% for maltose and 25% for tryptone and PYE medium).

### Both the major and alternate chemotaxis operons are involved in biofilm regulation

As chemotaxis is involved in surface colonization and biofilm formation in different microorganisms, our next step was to test populations of mutant strains were able to bind to surfaces as efficiently as WT populations. We first quantified the amount of biofilm formed after 24 hours under static conditions, using the five aforementioned carbon sources and PYE medium. Mutant phenotypes could be rescued by expressing a copy of the deleted gene on a replicating plasmid under control of a constitutive promoter (Fig. S2B).

No difference in adhesion could be detected in complex PYE medium, suggesting that neither chemotaxis operon is involved in long term biofilm formation when nutrients are not limiting (Fig. 3A-B). The Δ*cheAI* mutant was severely impaired in biofilm formation when grown in defined M2 medium, regardless of the tested carbon source. The deficiency ranged from 50-70% of WT levels, depending on the given carbon source (Fig. 3A-B). As CheAI is involved in chemotaxis (Fig. 2), we conclude that chemotaxis and biofilm formation pathways are interconnected through this crucial histidine kinase. Intriguingly, the Δ*cheBI* mutant formed more biofilm than WT when grown in defined media (Fig. 3A-B). This Δ*cheBI* mutant phenocopies the Δ*cheAI* mutant for chemotaxis but exhibits an inverse adhesion phenotype. These data suggest that, in addition to its known function as a methyltransferase modulating the chemotactic response, CheBI plays a role in biofilm regulation in *C. crescentus*. Interestingly, adhesion in the Δ*cheAII* mutant was reduced when grown with certain carbon sources; while biofilm formation of the Δ*cheAII* mutant was not significantly different from that of WT when grown with glucose, sucrose, or tryptone, it dropped to Δ*cheAI* levels in M2 supplemented with maltose or xylose (Fig. 3A-B). Thus, even if CheAII is not involved in chemotaxis (Fig. 2), this protein plays a role in biofilm regulation (Fig. 3A-B). This regulation is carbon source specific and is not dependent on chemotaxis. The Δ*cheBII* mutant phenocopied the Δ*cheBI* mutant and displayed a hyper-biofilm phenotype (Fig. 3A-B), suggesting that both CheB proteins are somehow involved in biofilm regulation. CheA and CheB proteins act in opposition; while mutations in *cheA* yield a decrease in adhesion, *cheB* deletions trigger an increase in adhesion.

**Figure 3:**
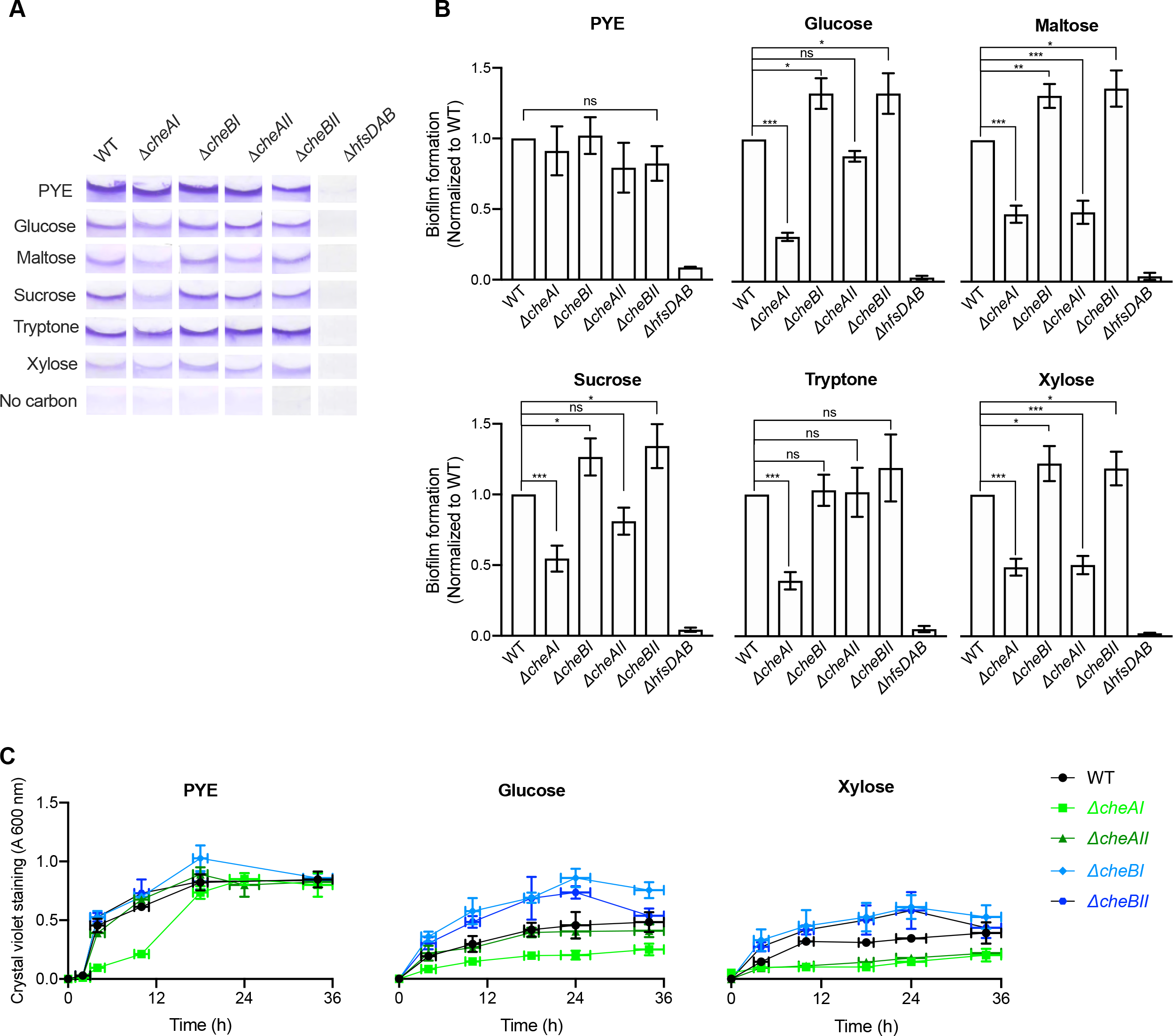
Biofilm formation. **(A)** Representative images of CV-stained biofilms grown on PVC coverslips under static conditions. **(B)** Biofilm formation after 24 h incubation at 30°C in PYE and M2 media with different carbon sources. Results are expressed as a percentage of WT biofilm formation in the given medium. Error bars represent the SEM of at least three independent replicates run in duplicate. Statistical comparisons were calculated using unpaired t-tests. *** *P* < 0.001; ** *P* < 0.01; * *P* < 0.1; ns = not significant.

To determine what stage of biofilm formation was impacted in these mutants, we monitored their attachment kinetics in the same static system. We focused on PYE and M2 supplemented with glucose (M2G) or xylose (M2X), since the adhesion phenotypes of the Δ*cheAI* and Δ*cheAII* mutants were different when grown in these different media. Indeed, in PYE, both Δ*cheAI* and Δ*cheAII* phenocopied the WT strain after a 24 hours incubation (Fig. 3B); however, when glucose was used as the sole carbon source, only Δ*cheAI* mutants were significantly impaired for biofilm formation after 24 hours (30% of biofilm formed compared to WT and Δ*cheAII*; Fig. 3B). Finally, in M2X, when xylose was the sole carbon source, both Δ*cheAI* and Δ*cheAII* mutants formed 50% less biofilm than WT (Fig. 3B). In PYE, when monitored over time, the amounts of biofilm formed by the Δ*cheAII*, Δ*cheBI*, and Δ*cheBII* mutants were similar to that of WT, highlighting that these proteins are not involved in biofilm regulation in complex medium. However, the Δ*cheAI* mutant was impaired in biofilm initiation, with only 30% of biomass attached compared to the WT strain after the first 10 hours (Fig. 3C). The biofilm formation of the Δ*cheAI* mutant eventually reached the same level as the WT strain after longer incubation times, showing that biofilm maturation is not impacted in this strain (Fig. 3B-C). In M2G and M2X, the Δ*cheAI* mutant was impaired in biofilm initiation, and this defect persisted through time, yielding around 30% of WT attached biofilm regardless of the incubation time (Fig. 3C). In both CheB mutants, the first steps of adhesion were more efficient than WT, with 3-5 times more cells attached to the surface within the first 4 hours of incubation (Fig. 3C). The attached biomass increased over time, and these mutants reached biofilm maturation earlier than did WT (Fig. 3C). The behavior of the Δ*cheAII* mutant was dependent on the carbon source; while it phenocopied WT in presence of glucose, it behaved like the Δ*cheAI* mutant in the presence of xylose (Fig. 3C).

Taken together, these results show an intriguing relationship between chemotaxis and biofilm regulation. While both CheAI and CheBI have been shown to be crucial for chemotaxis, these proteins have distinct antagonistic roles in biofilm regulation. In addition, these data highlight a role for the alternate chemotaxis operon in the regulation of biofilm formation through CheAII and CheBII. Furthermore, like their major chemotaxis operon counterparts, CheAII and CheBII also act in an opposite manner but do so conditionally, since CheAII is involved in biofilm regulation only in the presence of certain carbon sources.

### CheA and CheB proteins regulate holdfast production in an antagonistic manner

The adhesive holdfast is essential for long term adhesion in *C. crescentus* (29–31). Because long term adhesion was altered in the tested Che mutants, we sought to determine whether changes in holdfast production could be responsible for this defect. To quantify the proportion of single cells harboring a holdfast in the mixed population, we stained early exponential cultures grown in PYE, M2G, or M2X with fluorescent wheat germ agglutinin (WGA), which specifically stains the the N-acetylglucosamine residues present in the holdfast (34), and quantified the number of cells with a holdfast by fluorescence microscopy.

As previously reported, the number of cells harboring a holdfast is drastically different when the cells are grown in complex PYE medium compared to nutrient-defined media M2X (43) or M2G (44). Interestingly, there was also a difference in holdfast formation depending on the carbon source used in the growth medium; around 20% of WT cells harbored a holdfast when grown in presence of glucose, while less than 10% did in the presence of xylose (Fig. 4A).

**Figure 4:**
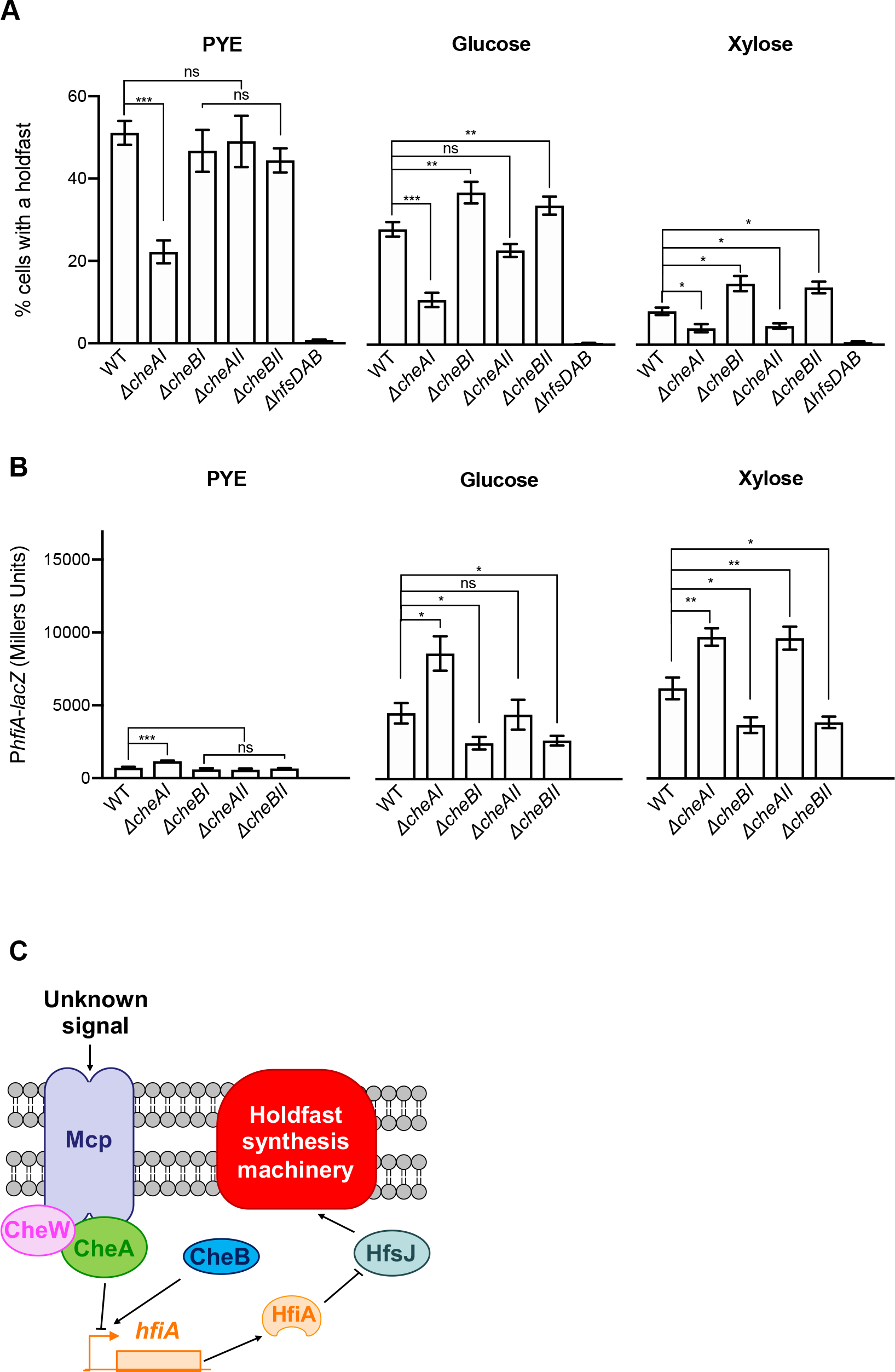
Role of the holdfast inhibitor HfiA. **(A)** Quantification of cells harboring a holdfast in mixed populations. Cells were stained using AF488-WGA and imaged by fluorescence microscopy. The results represent the average of three independent replicates (more than 300 cells per replicate) and the error bars represent the SEM. **(B)** ß-galactosidase activity of P*hfiA-lacZ* transcriptional fusions in PYE, M2G, and M2X media. The results represent the average of 9 independent cultures (assayed on 3 different days) and the error bars represent the SEM. Statistical comparisons were calculated using unpaired t-tests. *** *P* < 0.001; ** *P* < 0.01; * *P* < 0.1; ns = not significant. **(C)** Proposed model. Upon an unknown signal transduced via the MCP, CheA negatively regulates expression of the holdfast synthesis inhibitor gene, *hfiA*, leading to increased holdfast production. On the other hand, CheB promotes *hfiA* expression, resulting in a decrease in holdfast synthesis.

In all tested media, the number of cells producing a holdfast in the Δ*cheAI* mutant population was reduced by half compared to WT (Fig. 4A). The number of holdfasts detected in the other tested mutants did not significantly differ from the WT population (Fig. 4A). However, in defined media, both the Δ*cheB* mutant populations?? produced approximately 20% more holdfasts than WT (Fig. 4A). The number of holdfasts present in the Δ*cheAII* mutant phenocopied WT in M2G and the Δ*cheAI* strain in M2X (Fig. 4A). These results are in agreement with the biofilm phenotypes presented in Fig. 3.

### Transcription of the holdfast inhibitor-encoding gene, *hfiA*, is regulated by CheA and CheB

In *C. crescentus*, holdfast production is regulated by nutrient availability via the <u>h</u>old<u>f</u>ast <u>i</u>nhibitor protein, HfiA (43). In defined M2G and M2X media, *hfiA* expression is upregulated, resulting in a significant decrease in holdfast production compared to cells grown in PYE (43, 44). To determine if HfiA was playing a role in the differences in holdfast production observed in the present study, we measured the expression of *hfiA* using *lacZ* transcriptional fusions in WT and the chemotaxis mutants. In PYE, the activity of the *hfiA* promoter was minimal. Moreover, *hfiA* expression in the Δ*cheAI* mutant was approximately 50% higher than that in the other tested strains (Fig. 4B). In defined M2 media, we observed a drastic increase in *hfiA* transcription compared to in PYE as previously reported REF (Fig. 4B). There was also an overall increase in *hfiA* expression in M2X compared to M2G. In defined M2 media, *hfiA* expression was elevated in the Δ*cheAI* strain while it was decreased in both Δ*cheB* mutants (Fig. 4B). These results correlated with holdfast quantification (Fig. 4A); *hfiA* expression was lower in populations that have more cells with a holdfast. Overall, these observations suggest that chemotaxis genes regulate *hfiA* expression and thereby controls holdfast production in response to the carbon sources available in the medium (Fig. 4C).

## DISCUSSION

In this work, we investigated the putative roles of two chemotaxis operons in chemotaxis and surface attachment. We showed that only the major operon is involved in chemotaxis, while both operons regulate biofilm formation and holdfast production. Our results support a model where both CheAI and CheAII proteins negatively regulate the expression of the gene encoding the holdfast inhibitor protein, HfiA, while CheB proteins activate its expression in response to the carbon source present in the medium. In turn, this change in *hfiA* expression modulates holdfast production by interaction with the holdfast polysaccharide glycosyltransferase HfsJ (Fig. 4C). It has been recently shown that disturbances in flagellum or pilus synthesis modify holdfast production via *hfiA* regulation (44, 58), and we now add chemotaxis proteins as feedback regulators of holdfast synthesis via HfiA.

Other chemotaxis proteins, specifically, the CheY-like Cle proteins, have been shown to be involved in chemotaxis and holdfast synthesis (53). Interestingly, only CleA, located in the major chemotaxis operon, has been shown to play a role in chemotaxis regulation, confirming our observations that only the major chemotaxis operon properly functions as a regulator of chemotaxis (53). In addition, Cle proteins are involved in the regulation of holdfast production upon surface contact (53). However, holdfast synthesis by surface contact stimulation does not occur in defined M2 medium (44), suggesting that the regulation observed in our work is linked to developmentally programmed holdfast production and, therefore, occurs via a different mechanism. The multifunctional response regulator MrrA, which is essential for the general stress response in *C. crescentus*, has been recently shown to play a role in chemotaxis and holdfast production via the modulation of *hfiA* expression (59). It could be interesting to test if this global regulator is involved in holdfast regulation by the Che proteins. Another putative player to explore in the future is cdG. Indeed, this ubiquitous messenger molecule is involved in chemotaxis (53), holdfast production (40, 41), and *hfiA* expression (44), and it could be involved in the modulation of holdfast synthesis by the chemotaxis proteins.

Many studies in other bacteria have shown that chemotaxis-like operons are involved in behaviors independent from chemotaxis, such as cell differentiation or biofilm formation (3, 60). For example, regulatory pathways homologous to the chemotaxis system have been shown to control cyst formation in *Rhodospirillum centenum* (61), fibril polysaccharide production in *Myxococcus xanthus* (62), and cell aggregation and biofilm formation in *Azospirillum brasilense* (10). In these species, different proteins within the chemotaxis-like systems act antagonistically. In *R. centenum*, MCP, CheW, CheR, and CheA positively regulate cyst formation, while CheY and CheB are negative regulators (61). In *M. xanthus*, the chemotaxis-like Dif pathway regulates formation of fibril polysaccharide, a crucial component for fruiting body and spore formation. Within the Dif operon, DifD (CheY homolog) and DifG (CheC phosphatase homolog) are negative regulators of fibril production, whereas DifA (MCP homolog), DifC (CheW homolog), and DifE (CheA homolog) are positive regulators (62–64). In *A. brasilense*, CheA and CheY repress exopolysaccharide (EPS) production involved in cell-cell aggregation and biofilm formation, while CheB and CheR enhance it (10). Our results show that, in *C. crescentus*, different Che proteins also have antagonist effects on holdfast polysaccharide production, with CheA and CheB proteins from both operons acting as positive and negative regulators, respectively.

The best characterized chemotaxis-like operon involved in biofilm formation is the Wsp system of *Pseudomonas aeruginosa*. This system controls EPS production by modulating cdG levels in response to contact with a surface (65, 66). Briefly, WspR is a CheY homolog and a diguanylate cyclase that acts as the final response regulator of the Wsp system (65, 67). In its active form, WspR produces cdG, which in turn activates the production of Pel and Psl exopolysaccharides and biofilm formation. In that system, WspF (CheB homolog) is a modulator of WpsR. Deletion of *wpsF* results in increased phosphorylation of WspR, which negatively regulates the polysaccharides Psl and Pel while interfering with the intracellular levels of cdG (65, 67). Future work will determine if holdfast regulation by CheA / CheB in *C. crescentus* occurs using a similar mechanism.

In conclusion, we have demonstrated that the two chemotaxis operons of *C. crescentus* have distinct roles. Our data show that, while the major operon is involved in both chemotaxis and holdfast production, the alternate operon is a chemotaxis-like system involved in holdfast regulation but not chemotaxis to the compounds tested. Mutants lacking the response regulators CheAI or CheAII are impaired in cell attachment, presumably resulting from a defect in holdfast production, while CheBI and CheBII mutants produce more holdfasts and form more robust biofilms. We also showed that the regulation of holdfast synthesis by Che proteins is due at least in part to the modulation of the expression of *hfiA*, encoding the holdfast inhibitor protein HfiA. These data suggest a model where, in response to a yet unknown signal recognized by an MCP, CheAs promote *hfiA* expression and holdfast synthesis, while CheBs repress them (Fig. 4C). Further exploration of the mechanism by which this occurs may reveal how bacteria respond to external stimuli to optimize bacterial adhesion and surface colonization in various environments.

## MATERIALS AND METHODS

### Bacterial strains, plasmids, and growth conditions

The bacterial strains used in this study are listed in Table S2. *C. crescentus* strains were grown at 30°C in defined M2 medium (68) supplemented with 0.2% (w/v) of a given carbon source (listed in Table S1) or in complex peptone-yeast extract (PYE) medium (51). When appropriate, 5 μg/ml kanamycin or 1 μg/ml tetracycline was added to the medium. For induction of the Van promoter in pMT630-derivative constructs, 0.5 mM vanillate was added to the cultures before inoculation and incubation. *E. coli* Silver α select cells (Bioline) were used for cloning and were grown in LB medium at 37°C with 25 μg/ml kanamycin or 10 μg/ml tetracycline when appropriate.

In-frame deletion mutants were obtained by double homologous recombination, as previously described (69). Briefly, PCRs were performed using *C. crescentus* CB15 genomic DNA as the template to amplify 500-bp fragments from the upstream and downstream regions of the gene to be deleted. The primers designed for these in-frame deletions are listed in Table S3. PCR fragments were gel-purified and then digested by *Bam*HI and *Xho*I or *Xho*I and *Hind*III for upstream or downstream fragments, respectively. Purified, digested fragments were then cloned into the suicide vector pNPTS138 that had been digested by *Bam*HI and *Hind*III. The pNPTS138-based constructs were transformed into *E. coli* Silver α select cell and then introduced into *C. crescentus* by electroporation. The two-step recombination was carried out using sucrose resistance and kanamycin sensitivity (69). Then, the mutants were checked by sequencing to confirm the presence of the deletion.

The complementation plasmids, harboring *cheAI, cheAII, cheBI, or cheBII*, were constructed as follows. *C. crescentus* CB15 gDNA was used as template to PCR-amplify the genes of interest using primers containing *Hind*III (forward primers) and *Kpn*I (downstream primers) restriction sites (Table S2). PCR products were purified using the Qiaquick kit (Quiagen), digested using *Hind*III and *Kpn*I, and ligated into plasmid pMR10 (70) digested by the same enzymes. The plasmid expressing *pdeA* was constructed using the same method but using *Nde*I and *Kpn*I restrictions enzymes and the pMT630 plasmid (71).

### Growth curves and generation time calculations

Bacterial growth in the different media was measured in 3 ml liquid cultures (in 15 ml glass tubes) with shaking at 300 rpm. Overnight cultures were diluted in the same culture medium to an optical density (OD_600_) of 0.05 and incubated for 24 h. OD_600_ was measured at various time intervals to generate growth curves (OD_600_ vs time). Generation times were calculated from the exponential part of the growth curves, using the single exponential-growth function in the GraphPad Prism 6 software. Experiments were run in triplicates and all generation times were normalized to the average of WT doubling time in a given medium.

### Motility assays in semisolid media

Motility assays were performed using semisolid agar plates. Plates were poured using PYE or M2 medium supplemented with 0.2% (w/v) of the appropriate carbon source and 0.4% (w/v) noble agar (Difco). Cells were stabbed in the soft agar and incubated in a humid chamber at 30°C for 5 days. The diameter of the swimming ring formed by each tested strain was measured manually.

### Biofilm assays

Biofilm assays in multiwell plates were performed using two different setups that yield similar results (72): 1), adhesion to polyvinyl chloride (PVC) microscope coverslips placed vertically in plastic 12-well plates; or 2), adhesion to the inside surface of the wells of untreated, plastic 24-well plates. Bacteria were grown to mid-log phase (OD_600_ of 0.4 - 0.8) in the chosen medium and diluted to an OD_600_ of 0.05 in the same medium in 3 or 0.5 ml for the 12 or 24-well plate setup, respectively. Plates were incubated at 30°C for different times. Biofilms attached to coverslips or inside surfaces of the wells were quantified, as follows: wells or coverslips were rinsed with distilled H_2_O to remove non-attached bacteria, stained using 0.1% crystal violet (CV), and rinsed again with dH_2_O. The CV from the stained attached biomass was eluted using 10% (v/v) acetic acid and was quantified by measuring absorbance at 600 nm (A_600_). Biofilm formation was normalized to A_600_ / OD_600_ and expressed as a ratio of the WT level.

### Holdfast quantification using fluorescently labeled WGA lectin

The number of cells harboring a holdfast in mixed populations was quantified by fluorescent microscopy. Holdfasts were detected with AF488-WGA. Early exponential-phase cultures (OD_600_ of 0.2 − 0.4) were mixed with AF488-WGA (0.5 μg/ml final concentration). One microliter of WGA-stained cells was spotted on a 1.5-mm glass coverslip and covered with an agarose pad (1% SeaKem LE Agarose dissolved in dH_2_O). Holdfasts were imaged by epifluorescence microscopy using a Nikon Ti-2 microscope with a Plan Apo 60X objective, a GFP/DsRed filter cube, a Hamamatsu ORCA Flash 4.0 camera, and Nikon NIS Elements imaging software. The number of individual cells with a holdfast was calculated manually from microscopy images in the Nikon NIS Elements imaging software.

### β-galactosidase assays

Strains bearing transcriptional the reporter plasmid of the *hfiA* gene promoter fused to *lacZ* (44) were inoculated from freshly grown colonies into 5 ml of a chosen medium containing 1 μg/ml tetracycline and were then incubated at 30°C overnight. Cultures were then diluted in the same culture medium to an OD_600_ of 0.05 and incubated until an OD_600_ of 0.15 – 0.25 was reached. β-galactosidase activity was measured colorimetrically as described previously (73). A volume of 200 μl of culture mixed with 600 μl of Z buffer (60 mM Na_2_HPO_4_, 40 mM NaH_2_PO_4_, 10 mM KCl, 1 mM MgSO_4_, 50 mM ß-mercaptoethanol), 50 μl of chloroform, and 25 μl of 0.1% SDS. Two-hundred microliters of substrate *o*-nitrophenyl-β-D-galactoside (4 mg/ml) was then added to the cell mixture, and time was recorded until development of a yellow color. The reaction was stopped by adding 400μl of 1M Na_2_CO_3_ to raise the pH to 11. Absorbance at 420 nm (A_420_) was measured and the Miller Units of β-galactosidase activity were calculated as (A_420_ × 1000)/((OD_600_ × *t*) × *v*), where *t* is the incubation time in minutes and *v* is the volume of culture (in ml) used in the assay. The β-galactosidase activity of CB15 WT plac290 (empty vector control) was used as a blank sample reference.

## ACKNOWLEDGMENTS

The authors thank the members of the Brun laboratory and S. Zappa for critical reading of this manuscript. This study was supported by grant R35GM122556 from the National Institutes of Health and by a Canada 150 Research Chair in Bacterial Cell Biology to YVB.

---

**Figure S1:**
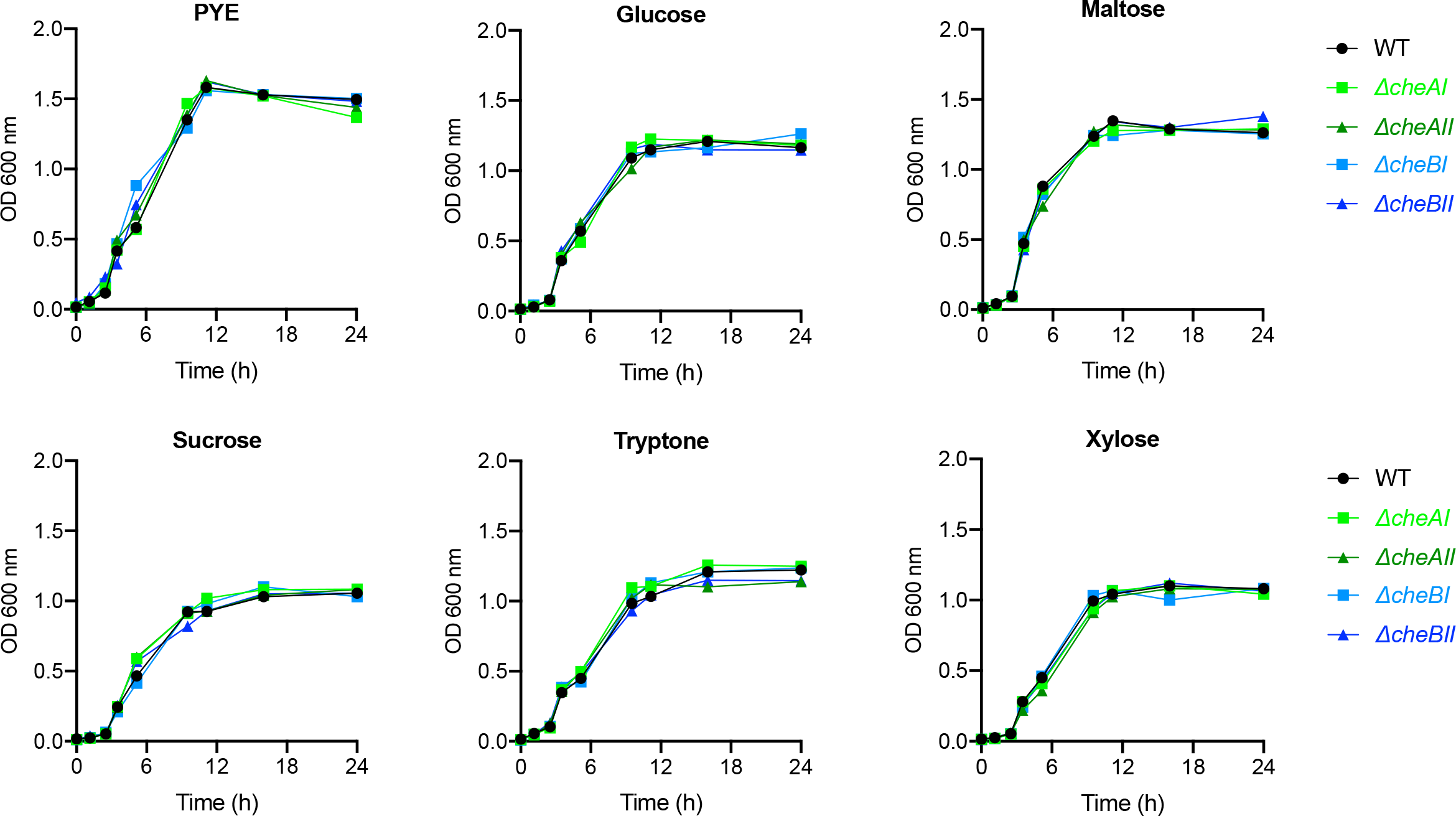
Growth in M2 with different carbon sources. Representative growth curves of cultures grown at 30°C and 300 rpm shaking, following OD_600_ over time. CB15 WT is shown in black circles, Δ*cheAI* and Δ*cheAII* in green squares and triangles, respectively, and *cheBI* and Δ*cheBII* in blue squares and triangles, respectively.

**Figure S2:**
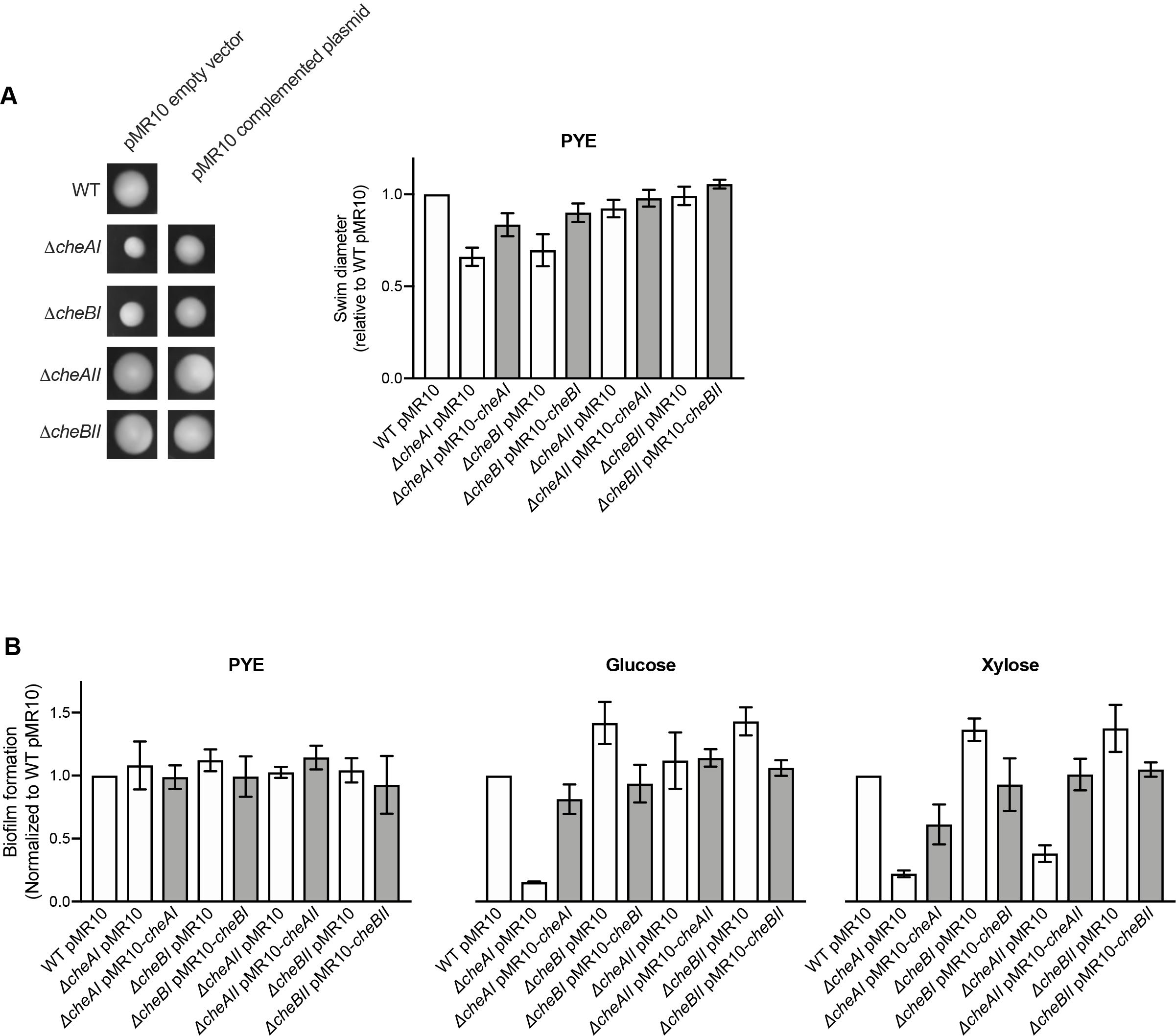
Phenotypes of Δ*cheA* and Δ*cheB* complemented strains. **(A)**Motility in semisolid agar PYE plates containing 0.4% noble agar and 5 μg/ml kanamycin. Results are expressed as the swim diameter relative to that of CB15 WT with pMR10 empty vector. **(B)**Biofilm formation in 24-well plates (after 24 h incubation at 30°C) in PYE, M2G, or M2X media supplemented with 5 μg/ml kanamycin. Results are expressed as a percentage of WT MR10 empty vector biofilm formation. Empty vector constructs and complemented strains are shown in white and gray bars, respectively. Error bars represent the SEM of three independent replicates, each run in duplicate.

**TABLE S1:**
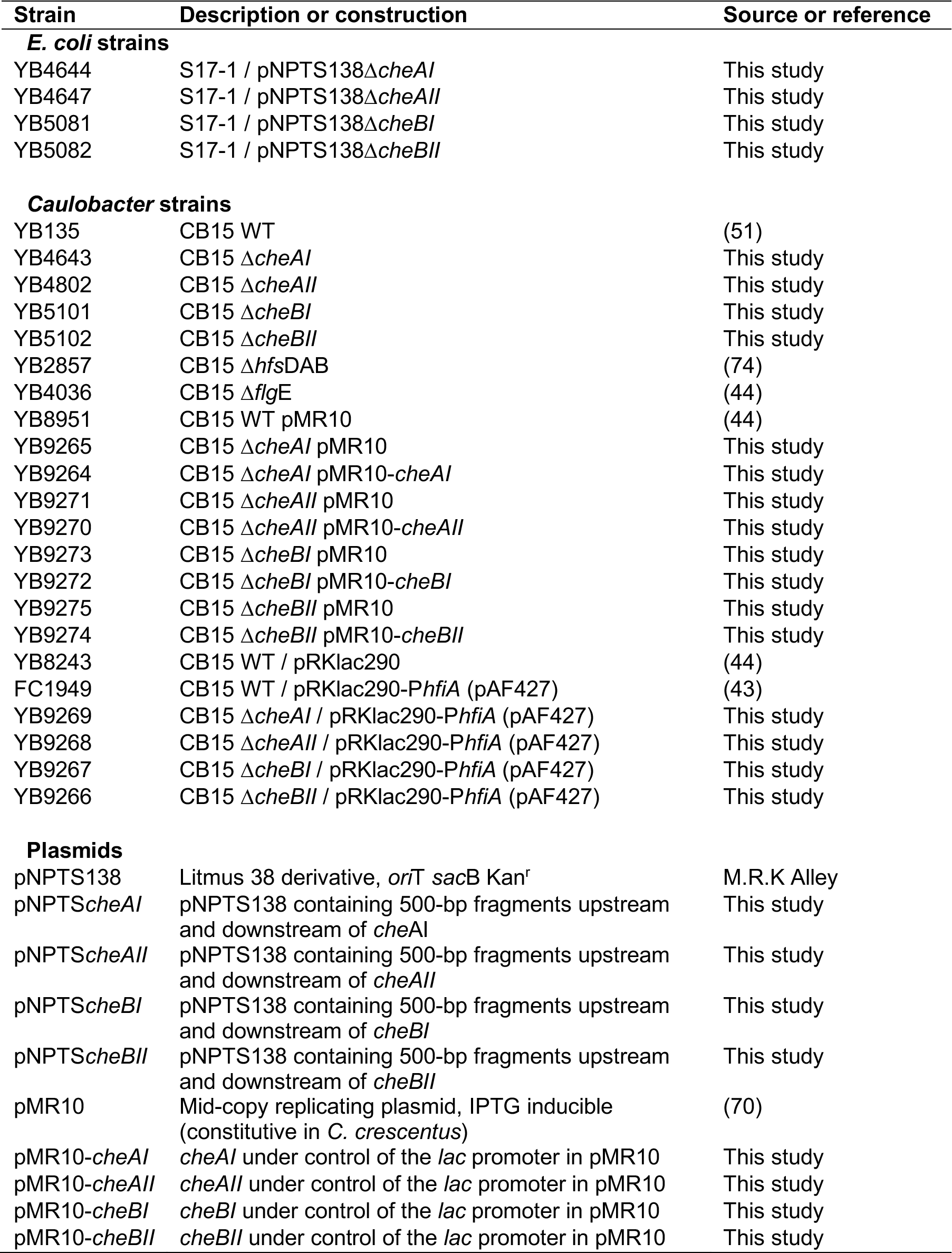
Bacterial strains and plasmids used in this study.

**TABLE S2:**
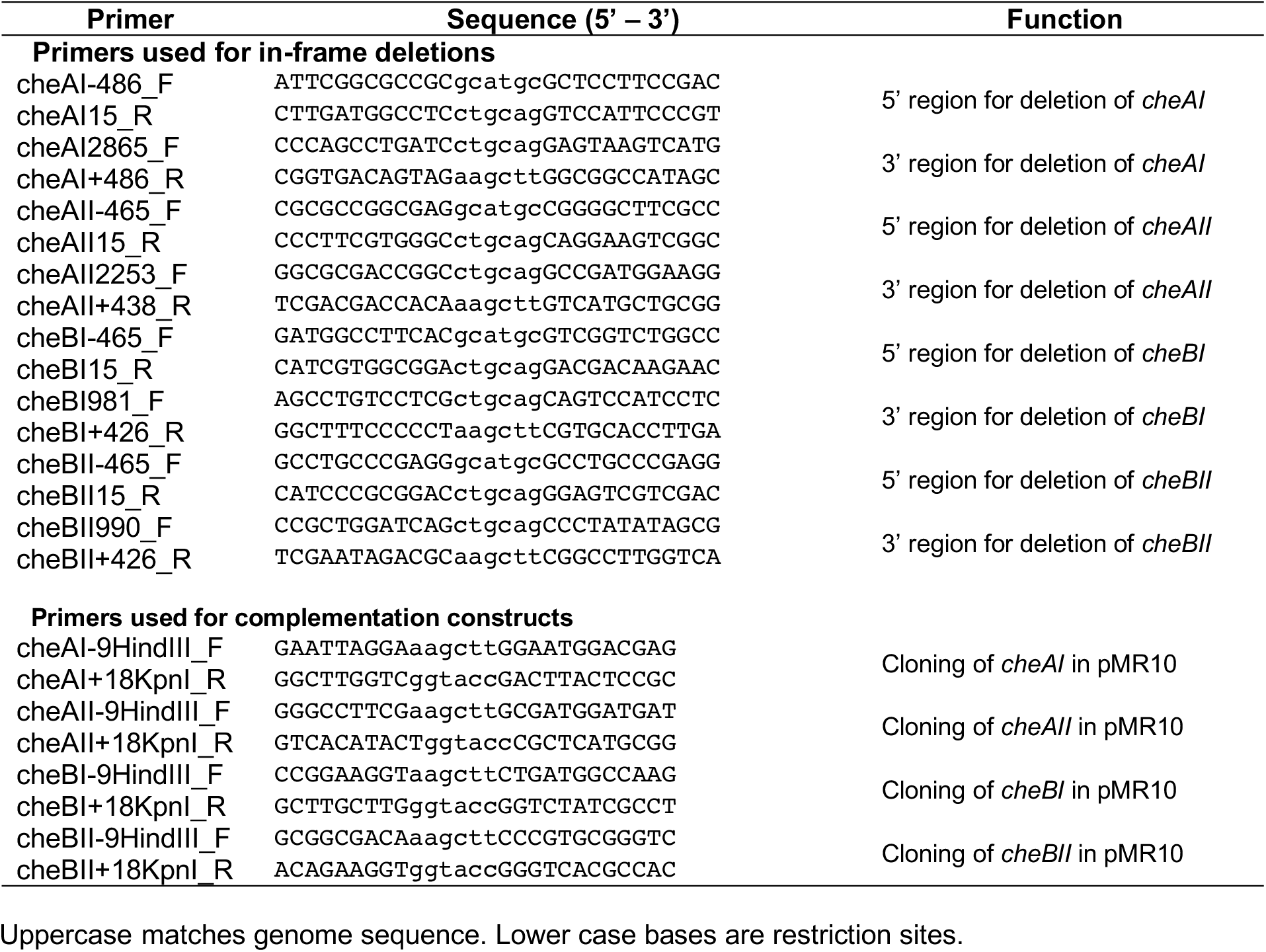
PCR primers used in this study.

